# Interface Refinement of Low-to-Medium Resolution Cryo-EM Complexes using HADDOCK2.4

**DOI:** 10.1101/2021.06.22.449462

**Authors:** Tim Neijenhuis, Siri C. van Keulen, Alexandre M.J.J. Bonvin

## Abstract

A wide range of cellular processes require the formation of multimeric protein complexes. The rise of cryo-electron microscopy (cryo-EM) has enabled the structural characterization of these protein assemblies. The produced density maps can, however, still suffer from limited resolution, impeding the process of resolving structures at atomic resolution. In order to solve this issue, monomers can be fitted into low-to-medium resolution maps. Unfortunately, the produced models frequently contain atomic clashes at the protein-protein interfaces (PPIs) as intermolecular interactions are typically not considered during monomer fitting. Here, we present a refinement approach based on HADDOCK2.4 to remove intermolecular clashes and optimize PPIs. A dataset of 14 cryo-EM complexes was used to test eight protocols. The best performing protocol, consisting of a semi-flexible simulated annealing refinement with restraints on the centroids of the monomers, was able to decrease intermolecular atomic clashes by 98% without significantly deteriorating the quality of the cryo-EM density fit.

## Introduction

Crucial processes in our cells such as metabolism, signal transduction, gene replication and transcription all involve an interplay between proteins. Understanding intermolecular interactions in complexes between proteins and other biomolecules is therefore key to obtain a deeper insight into these cellular mechanisms and their potential pathological effects (Arkin and Wells, 2004; Wells and McClendon, 2007). In order to analyze these multimeric protein complexes and their protein-protein interfaces (PPIs) high quality structures are required. Three common experimental methods to resolve protein structures are X-ray crystallography, nuclear magnetic resonance (NMR) and cryo-electron microscopy (cryo-EM), of which cryo-EM is the experimental method of choice to study large functional complexes (larger than ^~^80 kDa) (Bai et al., 2015; McPherson and Gavira, 2014). Since the development of direct electron detectors, cryo-EM has become widely used (Bai, McMullan and Scheres, 2015), which has resulted in a rapid growth of yearly deposited electron density maps (EDMs) (3827 in 2020) (*EMDB statistics*, 2021). The majority of the released maps originate from single particle (SPA) cryo-EM (77%), yielding an average resolution of 5.6 Å. Other cryo-EM techniques such as tomography, which allows *in-situ* detection, cannot yet reach such high resolutions. However, as the *ex-situ* conditions of SPA can affect a protein’s conformation, the lower resolution maps obtained with *in-situ* techniques are valuable for studying proteins under physiological conditions (Schur, 2019).

*De-novo* modeling of multimeric complexes based on lower quality electron density maps can be challenging. Often rigid-body fitting of available monomer structures (or homology models) into the EDM is used to build models of those protein complexes (Esquivel-Rodríguez and Kihara, 2013; Malhotra *et al*., 2019). Various software packages have been developed for this purpose, such as COAN (Volkmann and Hanein, 1999), DOCKEM (Roseman, 2000), EMFIT (Rossmann, 2000), Situs (Wriggers and Birmanns, 2001), UCSF Chimera (Pettersen *et al*., 2004), Flex-EM (Topf *et al*., 2008), BLC::EM-Fit (Woetzel *et al*., 2011) and PowerFit (van Zundert and Bonvin, 2015). After determining the initial position of each monomer, the models are often refined with respect to the EDM. An example of such an approach is the refinement of protein models in reciprocal space (e.g. by REFMAC) (Brown *et al*., 2015; Kovalevskiy *et al*., 2018). This technique allows the use of well-established methods developed for X-ray crystallography. Other common tools, like phenix.real_space_refine (Adams *et al*., 2010; Afonine *et al*., 2018) and Rosetta (Wang *et al*., 2016), use real space refinement to optimize models.

Despite the different complex refinement methods, low resolution cryo-EM models deposited in the protein databank (PDB) often contain a relatively high number of atomic clashes at the PPIs compared to higher resolution datasets such as the Docking Benchmark 5 (BM5) (Vreven *et al*., 2015) which includes a low average number of 0.38 ± 0.86 clashes at the interface. These clashes in cryo-EM models are artefacts of the rigid body fitting procedure and are often not resolved by refinement as electrostatics and stereochemical properties might not be fully enforced (Igaev *et al*., 2019). These clashes contribute to an incorrect or blurry representation of the interfaces in those protein complexes, making it challenging to extract information for further experiments (e.g. mutagenesis) or drug design.

Here, we present a refinement protocol to remove clashes from the PPI of low-to-medium resolution cryo-EM structures using our integrative modelling software package HADDOCK (Dominguez, Boelens and Bonvin, 2003), which has recently been extended to include up to 20 separate molecules per protein complex (Karaca *et al*., 2017). Eight different protocols are investigated in this study which are benchmarked on a set of 14 multimeric cryo-EM complexes with resolutions ranging from 5.1 Å to 9.3 Å. The results show that clashes at the interface of multimeric cryo-EM complexes can be reduced by up to 98% by applying a semi-flexible simulated annealing refinement with restraints on the centroids of the components of the complex without significantly compromising the fit to the EDM.

## Results

The 14 complexes in our benchmark (Figure 1, Table 1) were refined using our integrative modelling software package HADDOCK, version 2.4. In all cases the rigid body docking stage was skipped and the complexes were kept in their original orientation. For all protocols, the number of models to be generated was set to 50 for all stages of HADDOCK. The first five tested protocols use the semi-flexible simulated annealing (SA) stage of HADDOCK (it1) to refine the interfaces using various combinations of restraints (see Methods). We also tested the final refinement stage in explicit solvent (itw) as a stand-alone protocol. The last two protocols make use of the recent update of HADDOCK supporting coarse graining (CG) (Honorato, Roel-Touris and Bonvin, 2019; Roel-Touris *et al*., 2019). The last step of this CG docking protocol is the rebuilding of an all-atom (AA) model of the complex. This final CG-to-AA conversion step can also be used as a refinement protocol to remove intermolecular clashes. The 8 different protocols are summarized in Table 2 and details are provided in the Methods. The performance of the protocols was compared by analyzing the top 4 models ranked by HADDOCK.

**Figure 1.**
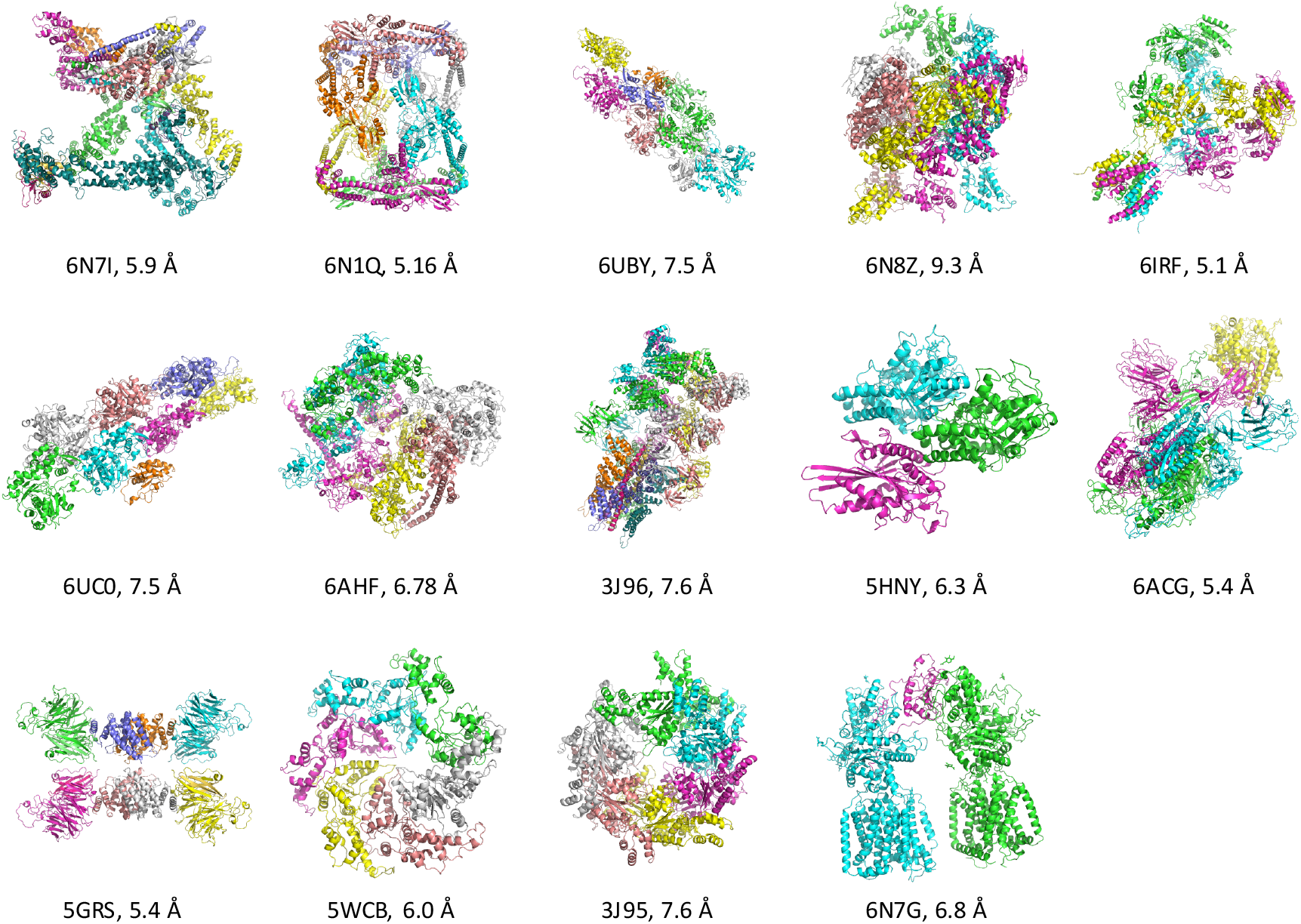
Cartoon representation, PDB IDs and cryo-EM map resolution of the selected 14 cryo-EM complexes. Each protein chain is highlighted by a unique color.

**Table 1.**
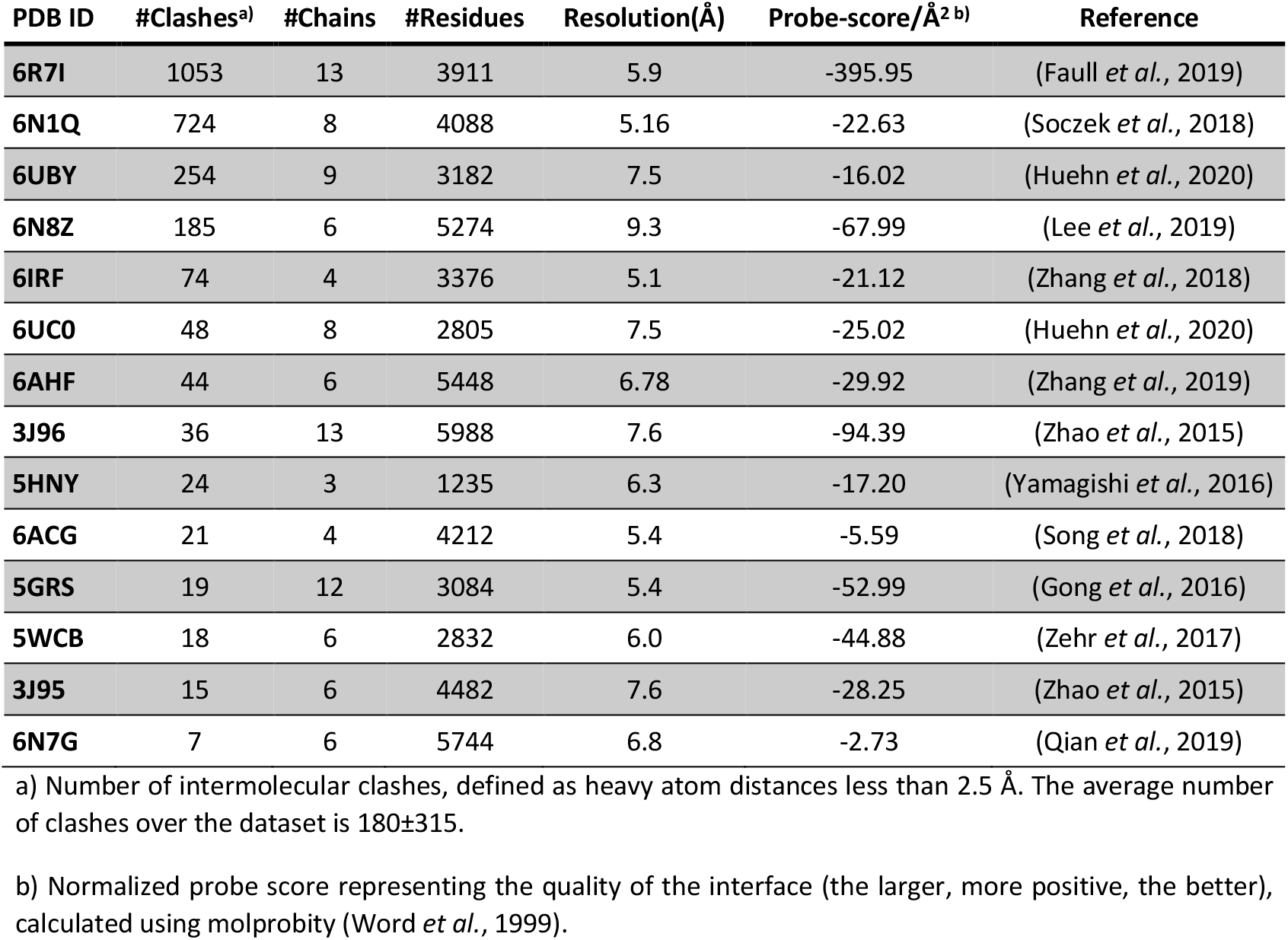
Dataset used for interface refinement ordered by number of atomic clashes at the interface.

**Table 2:** Overview of the tested refinement protocols.

| Protocol | Method (HADDOCK stage) | Additional restraints | ID |
| --- | --- | --- | --- |
| 1 | Semi-flexible simulated annealing (it1) | - | SA |
| 2 | Semi-flexible simulated annealing (it1) | Cα-Cα Distances | SA_CA |
| 3 | Semi-flexible simulated annealing (it1) | Surface restraints | SA_SR |
| 4 | Semi-flexible simulated annealing (it1) | Center of mass | SA_CM |
| 5 | Semi-flexible simulated annealing (it1) | Centroids | SA_CTRD |
| 6 | Solvated molecular dynamics (itw) | - | MD |
| 7 | Coarse grained back mapping | - | CG |
| 8 | Coarse grained back mapping and<br>Solvated molecular dynamics | - | CG_MD |

### HADDOCK Refinement Leads to Significant PPI Clash Removal

The first metric we used to assess the performance of the tested refinement protocols was the reduction in the number of interatomic clashes at the protein-protein interface. Overall, we observe a significant reduction in the number of clashes with respect to the reference starting structures. All protocols in which simulated annealing was used show a similar decrease in the number of clashes, from an average of 180 ± 315 in the deposited models (Table 1) to 4 ± 9 in the refined models, corresponding to an average reduction of 98.2% (Figure 2, 3A). When only a final refinement in explicit solvent was applied to the reference structures (protocol 6: MD), less clashes can be removed from the interface. We also observed an increase of interface clashes (from 6 in the reference structure to 27 after MD) for 6N7G, the complex with the lowest number of clashes. The average clash reduction is 63.1%, which is significantly lower than for the annealing protocols. A similar trend is observed for the CG-to-AA conversion refinement protocols in which the addition of a final refinement in explicit solvent (protocol 8: CG_MD) leads to a smaller reduction of intermolecular atomic clashes compared to CG-to-AA conversion alone (protocol 7: CG) (Figure 3A).

**Figure 2.**
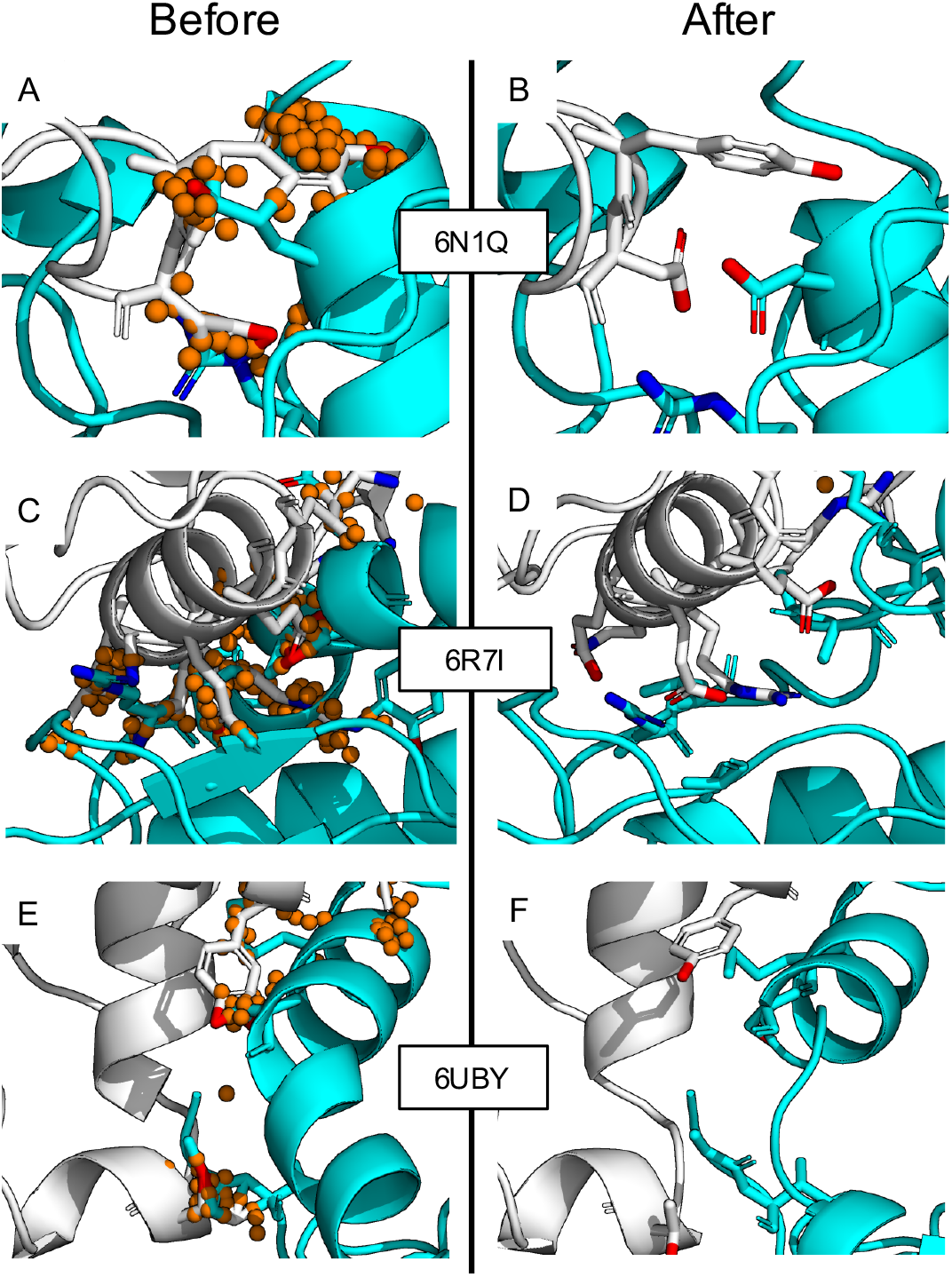
Illustration of intermolecular interactions for interfaces of three presentative complexes (6N1Q, 6R7I and 6UBY) before **(A,C,E)** and after **(B,D,F)** clash removal with HADDOCK using protocol 5, the simulated annealing refinement with centroid restraints (SA_CTRD). The orange spheres represent intermolecular clashes (atoms with intermolecular contacts < 2.5 Å). Unique protein chains are shown in cyan and gray.

**Figure 3.**
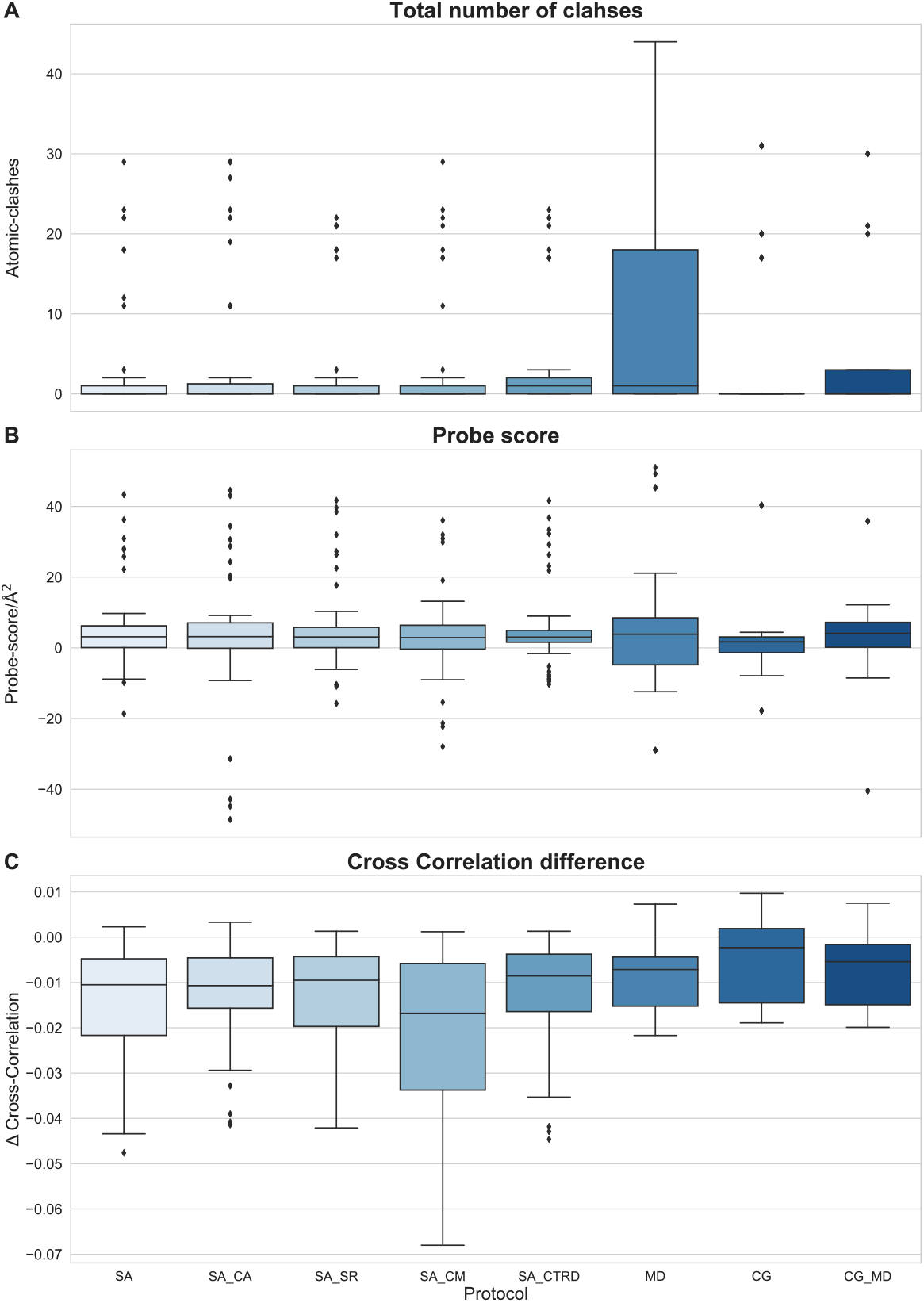
Atomic clashes, probe score, and delta cross-correlation per protocol (Table 2). Each box plot includes the top 4 ranking refined models of all 14 complexes for the corresponding protocol. (A) Number of remaining clashes. (B) Probe-score/Å2. (C) Delta cross-correlation with respect to the reference model.

### CG protocols Lead to the Best Cross-Correlation with the EM Density

Besides enhancing the quality of the interfaces, the overlap between the cryo-EM electron density map and the refined model also has to be preserved in order for the models to be in line with the experimental data. For each protocol, we calculated the cross-correlation between the refined model and the EM density (CC) and its change with respect to the starting reference structure (Δ_cc_) (Figure 3B). A positive Δ_cc_ indicates an increase in correlation (improvement of the fit with respect to the EM density) while a negative value indicates a decrease.

Generally, a slight decrease in cross-correlation was observed for all tested protocols (note that the cryo-EM density was not included as a restraint in the refinement). The different types of restraints in the simulated annealing protocols do not result in significantly different Δ_cc_ values, all around −0.01, except for CM restraints, which lead to an average Δ_cc_ of −0.02 ± 0.02. The MD protocol, which includes a picosecond time-scale MD simulation in explicit solvent, results in a similar Δ_cc_ compared to the simulated annealing protocols. The CG based protocols maintained the best correlation, with Δ_cc_ of −0.005 ± 0.009 and −0.007 ± 0.008, for CG and CG_MD respectively.

When taking into account only the analysis of the clash removal and density cross-correlation, we can conclude that the CG-to-AA conversion seems to be the best performing refinement method. Additionally, CG also outperforms the other protocols regarding computational load, requiring the least amount of CPU time of all protocols (Supplemental Figure 1).

### Interface Contact Quality is Enhanced by HADDOCK Refinement

Next, we evaluated the quality of the intermolecular contacts at the PPIs using Probe, which produces an interface score per Å^2^ (the more positive the score the better the quality of the interface) (Word *et al*., 1999). The average calculated probe score for the 14 reference complexes of the benchmark was found to be −60 ± 100 (Table 1). All of the tested refinement protocols were able to improve the probe score significantly compared to the reference structures, with average scores ranging between 1.85 and 5.5 (Figure 3C). In line with the clash removal analysis, the protocols including SA produce similar probe scores. Including surface (SA_SR) or centroid (SA_CTRD) restraints, results in an average score of 5.5, which slightly outperforms the base protocol SA with an average score of 5.3.

Despite being among the top performing protocols in terms of clash removal and density cross-correlation, interface refinement using CG-to-AA conversion, with (CG_MD) and without (CG) the final water refinement led to an average probe score of 1.8, which is significantly better than the reference complex structures, but ranks lowest compared to the other protocols.

### The Structural Quality of Protein Complexes is Best Preserved in All-Atom Protocols

Since the produced refined structures should maintain their structural integrity while improving the interface quality, we also analyzed the overall structural quality of the produced models. First, we calculated the Ramachandran and rotamer outliers for each model. Delta values with respect to the reference structures were obtained and are shown in Supplemental Figure 2A. An increase of Ramachandran outliers, close to 1%, was found for SA-based protocols while outliers remain close to 0 for the MD protocol. The largest number of outliers was detected in the CG protocols (CG and CG_MD).

Concerning the rotamer outliers, we observe an increase of more than 11% on average in all five SA protocols (Supplemental Figure 2B). The rotamer outliers show a similar trend with respect to the Ramachandran outliers for protocol MD, as less rotamer outliers are introduced in comparison to the simulated annealing protocols. Protocol CG has a similar performance as protocol MD. Protocol CG_MD, however, performs worst of all, with an average increase in rotamer outliers of 14%.

For all protocols except the CG refinement, we observe similar secondary structure conservation (Supplemental Figure 3) with respect to the reference structures. Models from protocol CG include on average a 10% decrease in secondary structure towards the more unstructured turns. Loss of helical structure contributes the most to this secondary structure shift (approximately 9%). The addition of MD simulations (CG_MD) enhances the number of secondary structure elements. Nonetheless, the percentage of strands and helices in protocol CG_MD remains significantly lower than for all SA and MD protocols.

It seems that clash removal by CG-to-AA conversion results in perturbations of the secondary structure, leading to protein models with larger regions that cannot be assigned as standard helix or strand. While the CG-to-AA back-mapping protocol performs best in clash removal and in maintaining the correlation to the experimental density, it does induce shifts in secondary structure towards less-structured turns conformations.

## Discussion

In order to establish which interface refinement approach of low-to-medium resolution cryo-EM structures leads to the highest quality complex structures, we have compared intermolecular clash removal, interface quality, density cross-correlation and secondary structure conservation. Because no single protocol scored best for all evaluated features, we have assessed the impact of each metric on the overall quality of the refined models to determine the best protocol. All criteria considered, the simulated annealing protocols (1-5) are the best performing in terms of model quality and secondary structure conservation. The top protocol, protocol 5 (SA_CTRD), including centroid restraints, requires minimal prior data generation and leads to enhanced results in density cross-correlation and interface quality, while resulting in similar clash removal compared to the other protocols. Additionally, no secondary structure perturbation is observed with respect to the reference, unrefined structures.

Protocol MD, including solely a short final refinement via an MD simulation in explicit solvent, performs similarly to protocol SA_CTRD in the delta cross-correlation and secondary structure analysis, and even outperforms protocol SA_CTRD in the rotamer and Ramachandran outliers. However, models refined by MD show the highest number of remaining intermolecular clashes with an average of 8.09 (±13.5). Therefore, this protocol is the least favorable in terms of clash removal, even if on average it decreased the number of clashes by 63.1%. Protocol CG, using CG-to-AA conversion, results in better crosscorrelation and PPI clash removal compared to protocol SA_CTRD. CG also shows an increase of cross correlation for several refined complexes. However, CG also leads to significant secondary structure perturbations (10%). One could argue that secondary structure assignment can be affected by the cutoff used to define a secondary structure element in the employed software. Yet, through manual inspection of the individual complexes, we have observed severe backbone distortions in CG produced complex structures (data not shown), suggesting this decrease in secondary structure is not solely a result of the cutoff definition but the result of the effective clash removal. Hence, we deem the CG protocols (protocol CG and CG_MD) less fit to yield accurate interface representations than the other protocols.

From all protocols tested, the SA refinement protocol with centroid restraints (protocol 5, SA_CTRD) appears to be the most optimal to produce realistic PPIs for large protein complexes (Supplemental Methods).

## Conclusion

In this study we have shown the potential of our integrative docking software HADDOCK for refining the protein-protein interfaces of low-to-medium resolution (5-10 Å) cryo-EM structures of large assemblies. The current version of HADDOCK (version 2.4) allows refinement of complexes including up to 20 subunits, which makes it especially suitable for structural refinement of large multimeric complexes.

After comparing all tested protocols, the protocol including a semi-flexible simulated annealing stage in combination with centroid restraints (SA_CTRD) appears to produce the highest quality complex structures, removing interatomic clashes by 98% and maintaining the secondary structure of the reference structure. Overall, the models resulting from this refinement approach provide a more realistic representation of the intermolecular interactions than the reference structures, which will facilitate the use of these models to address details of the recognition process and generate new hypotheses such as for residue mutation studies. This refinement protocol has now been implemented in the refinement interface of the HADDOCK2.4 web portal (https://wenmr.science.uu.nl/haddock2.4/refinement), which should facilitate its use.

## Methods

### Key Resource Table

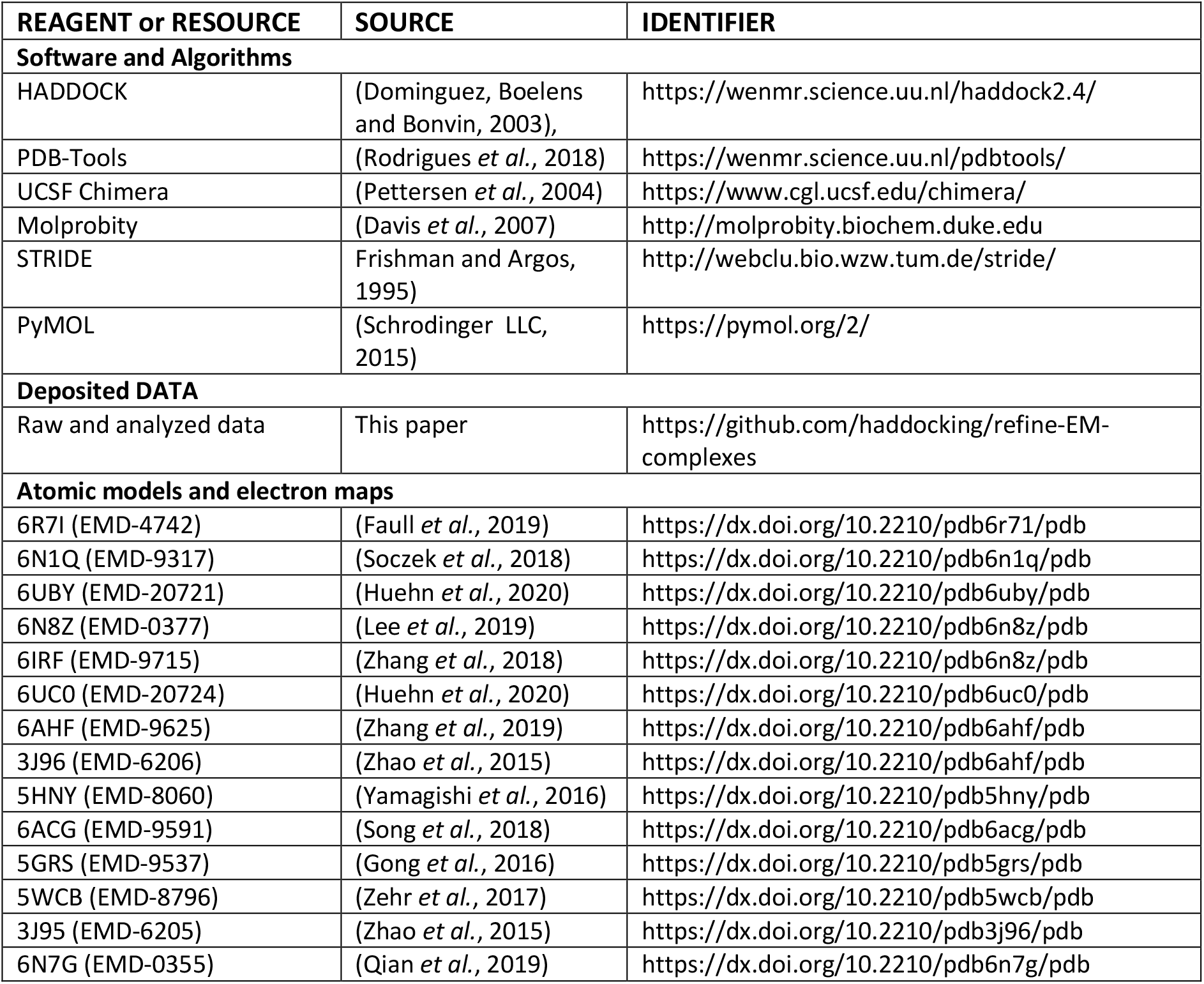

#### Dataset Generation and Preparation

The dataset used to benchmark the protocols described in this work is composed of 14 cryo-EM complexes in a resolution range of 5-10 Å. Complexes were selected based on the following criteria: (i) a chain count of up to 20, (ii) absence of nucleic acids and (iii) minimal redundancy within the dataset. The fourteen complexes selected are summarized in Table 1 and visualized in Figure 1. The structures were prepared using pdb-tools (Rodrigues *et al*., 2018). Co-factors distant from any interface were removed for convenience using `pdb_delresname.py` and noncanonical amino acids were substituted for their canonical counterparts using `pdb_rplresname.py` (e.g. MSE to MET). The individual chains were finalized for use in HADDOCK using `pdb_tidy.py`.

#### Refinement protocols

We used our integrative modelling software package HADDOCK, version 2.4, to refine the protein complexes in the dataset, testing several protocols to remove atomic clashes at the interfaces of these multimeric protein complexes. The default HADDOCK protocol consists of three stages. First, rigid body docking (it0) is performed in which each protein is treated as a rigid entity (Dominguez, Boelens and Bonvin, 2003). This docking step is usually guided by experimental or predicted interaction information, translated into distance restraints which bias specific protein regions to interact without predefining their relative orientation. Subsequently, the best models from it0 (200 by default) move to stage two (it1), which refines the PPIs via simulated annealing protocol in torsion angle space. Here, flexibility is gradually added to the protein interface, first for side chains and then both side chains and backbone, to allow for small conformational changes and optimize the intermolecular interactions. The third and last stage consists of a final refinement (itw). In this step, the protein-protein interface is refined by either an energy minimization (default in HADDOCK2.4) or by a very short molecular dynamics simulation using an explicit solvent shell. Eight different refinement protocols were considered in this work (Table 2). In all cases the rigid body docking stage (it0) was skipped and the complexes were kept in their original orientation (randomization of starting orientations turned off). For all protocols, the number of models to be generated was set to 50 for all stages of HADDOCK.

##### Semi-Flexible Interface Refinement (protocols 1-5)

The first five protocols use the semi-flexible simulated annealing stage of HADDOCK (it1) to refine the interfaces. The simplest form of this protocol, *protocol 1*, includes a simulated annealing step in which no additional restraints are provided to guide the refinement, followed by a final energy minimization in itw. For *protocol 2*, intermolecular Cα-Cα atom distances between residues at the PPI within an 8 Å cutoff are selected as restraints. To allow for movement of the individual chains, the upper distance limit is defined as the measured Cα-Cα distance padded with 1.5 Å. The lower bound distance was set to 0 Å. During refinement, 50% of the provided distance restraints is randomly excluded for each model generated. In *protocol 3*, “ambiguous surface restraints” are taken into account during it1. These are automatically calculated by HADDOCK as one ambiguous distance restraint per pair of components, defined between all Cα atoms of the two components with sum averaging and an upper limit of 7 Å. This restraint effectively enforces the two molecules to remain in contact while allowing for possible changes in the interfaces. The final restraints tested with the simulated annealing protocol are center of mass (CM) and centroid restraints in protocols 4 and 5, respectively. The center-of-mass restraints are automatically defined by HADDOCK between each pair of components as a distance restraint between the geometric center of the Cα atoms of the two components (center averaging). The upper distance limit for the restraints is the sum of the “effective radius” of each molecule, which is defined as half the average length of the two smallest principal components. Centroid restraints are distance restraints that effectively restrain the position of the geometric center of each component based on its Cα atoms to its original position.

##### Refinement in Explicit Water (protocol 6)

Besides protocols based on the it1 semi-flexible refinement, we have also tested the final refinement stage in explicit solvent (itw) as a stand-alone protocol (i.e. bypassing the it0 and it1 stage). In this setup (*protocol 6*), we used the experimental structure of the protein complex as a starting structure for itw. A solvent shell is built around the complex and, subsequently, a series of short MD simulations are performed using an integration time step of 2 fs: First, the system is heated to 300 K (500 MD steps at 100, 200 and 300 K) while all atoms except the side-chain atoms at the interface are restrained to their original position (force constant 5 kcal mol^−1^). Next, 1250 MD steps are performed at 300 K with position restraints (force constant 1 kcal mol^−1^) for heavy atoms which are not part of the PPI (residues not involved in intermolecular contacts within 5 Å). Finally, the system is cooled down (1000 MD steps at 300, 200 and 100 K) with position restraints on the backbone atoms of the protein complex, excluding the interface atoms (Dominguez, Boelens and Bonvin, 2003; Van Zundert *et al*., 2016).

##### Refinement by Coarse Graining (protocols 7 and 8)

The most recent update of HADDOCK supports coarse graining (CG) (Honorato, Roel-Touris and Bonvin, 2019; Roel-Touris *et al*., 2019) in which models can be coarse grained using the Martini force field (Monticelli *et al*., 2008) to reduce computational time as well as to smoothen the energy landscape. The last step of the CG docking protocol in HADDOCK is the rebuilding of an all-atom (AA) model of the complex. This final CG-to-AA conversion step can also be used as a refinement protocol to remove intermolecular clashes as recently demonstrated in a membrane complexes docking protocol combining rigid-body docking with Lightdock (Jiménez-García *et al*., 2018) and refinement in HADDOCK (Roel-Touris, Jiménez-García and Bonvin, 2020). CG-to-AA conversion is achieved by defining distance restraints between the atomic particles and the CG particle of which they are a part of. Using these distance restraints, the initial atomistic models are fitted onto the CG representation and the interfaces are optimized (for details see Roel-Touris et al., 2019).

For this protocol, we first converted the reference structures to coarse-grained models and generated the CG-to-AA restraints (automatically done on the web server). Then, it0 and it1 were disabled and only itw was used, which includes the CG-to-AA conversion. The tested CG protocols included the CG-to-AA conversion and a final energy minimization for protocol 7 or CG-to-AA conversion with a final MD refinement in explicit solvent (as done for protocol 6) for protocol 8.

#### Analysis

The performance of each protocol using the top 4 models of each complex according to their default itw HADDOCK score was assessed by analysing the intermolecular atomic clashes (i), the interface quality (ii), the cross-correlation with the cryo-EM density (iii), the overall structural quality (iv) and the secondary structure content (v). The first property we examined was the removal of atomic interface clashes (i) with respect to the reference structures. Atomic clashes in the protein complexes were calculated as intermolecular heavy atom contacts < 2.5 Å. The quality of the interface (ii) based on the contacts between chains was evaluated by using Probe (Word *et al*., 1999) in default mode. Calculated probe scores were normalized according to the surface area (Å^2^).

Besides the PPI quality, we also analysed the overlap between the experimental electron density map (EDM) and the models. For this the cross-correlation (iii) of each model with its experimental EDM was obtained by using the Chimera *fit in map* function (Pettersen *et al*., 2004). First, a density map of the atomic refined model was generated by describing each atom as a gaussian distribution with a width proportional to the resolution of the experimental map and an amplitude proportional to the atomic number. Subsequently, the maps were fitted iteratively (2000 steps), sampling over the three rotational and three translational axes until the overlap converged. The correlation was then calculated using equation 1, where *u* represents grid point values of the generated map and *v* the trilinear interpolated point values of the experimental map.

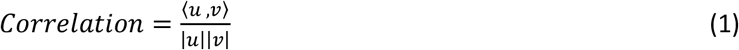

The overall structural quality (iv) of the models was checked by calculating the Ramachandran and rotamer outliers as well as the clash score of the whole complex via molprobity (Davis *et al*., 2007; Hintze *et al*., 2016; Williams *et al*., 2018). Finally, we calculated the percentage of secondary structure elements (v) of the protein complexes using STRIDE (Frishman and Argos, 1995) and compared the starting reference structures to determine the loss or gain in secondary structure.

Visualization of the complexes and preparation of the figures was performed using PyMOL (Schrodinger LLC, 2015).

## Data availability

The dataset used for this research, the top 4 refined HADDOCK models, raw data and analysis scripts are available at https://github.com/haddocking/refine-EM-complexes.

## Acknowledgments

This work has been performed with the financial support of the Dutch Foundation for Scientific Research (NWO) (PPS Technology Area grant 741.018.201)) and the European Union Horizon 2020 project BioExcel (823830).

