## Supplementary Material for "Interface Refinement of Low-to-Medium Resolution Cryo-EM Complexes using HADDOCK2.4"

### SUPPLEMENTAL FIGURES

**Supplemental figure 1.** Boxplots representing the CPU time used per refinement condition to calculate 1 model for each structure in the dataset.

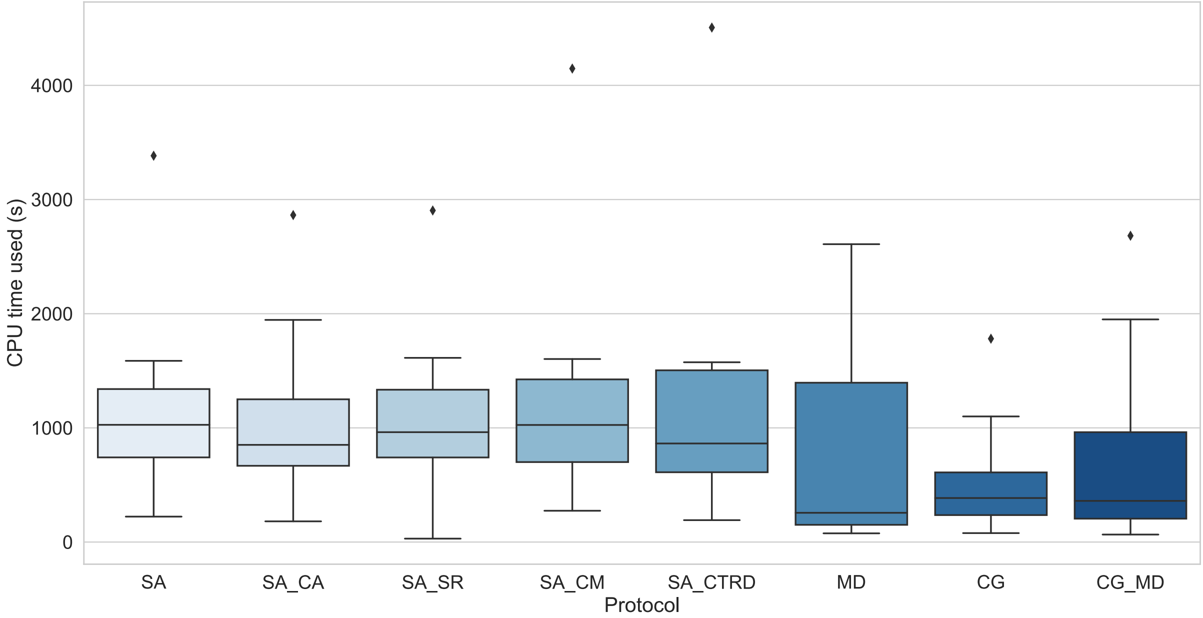

**Supplemental Figure 2.** Molprobity structural quality analysis results. The box plots represent delta values calculated by subtracting the reference from the model (model-reference) and are based on the top 4 refined models of all 14 complexes. **(A)** Delta Ramachandran outliers are shown with respect to the reference structures (negative values indicate an improvement with respect to the reference). **(B)** Delta rotamer outliers are shown with respect to the reference structures (negative values indicate an improvement with respect to the reference). **(C)** Delta clash scores (number of serious clashes for every 1000 atoms) are shown with respect to the reference structures (negative values indicate an improvement with respect to the reference).

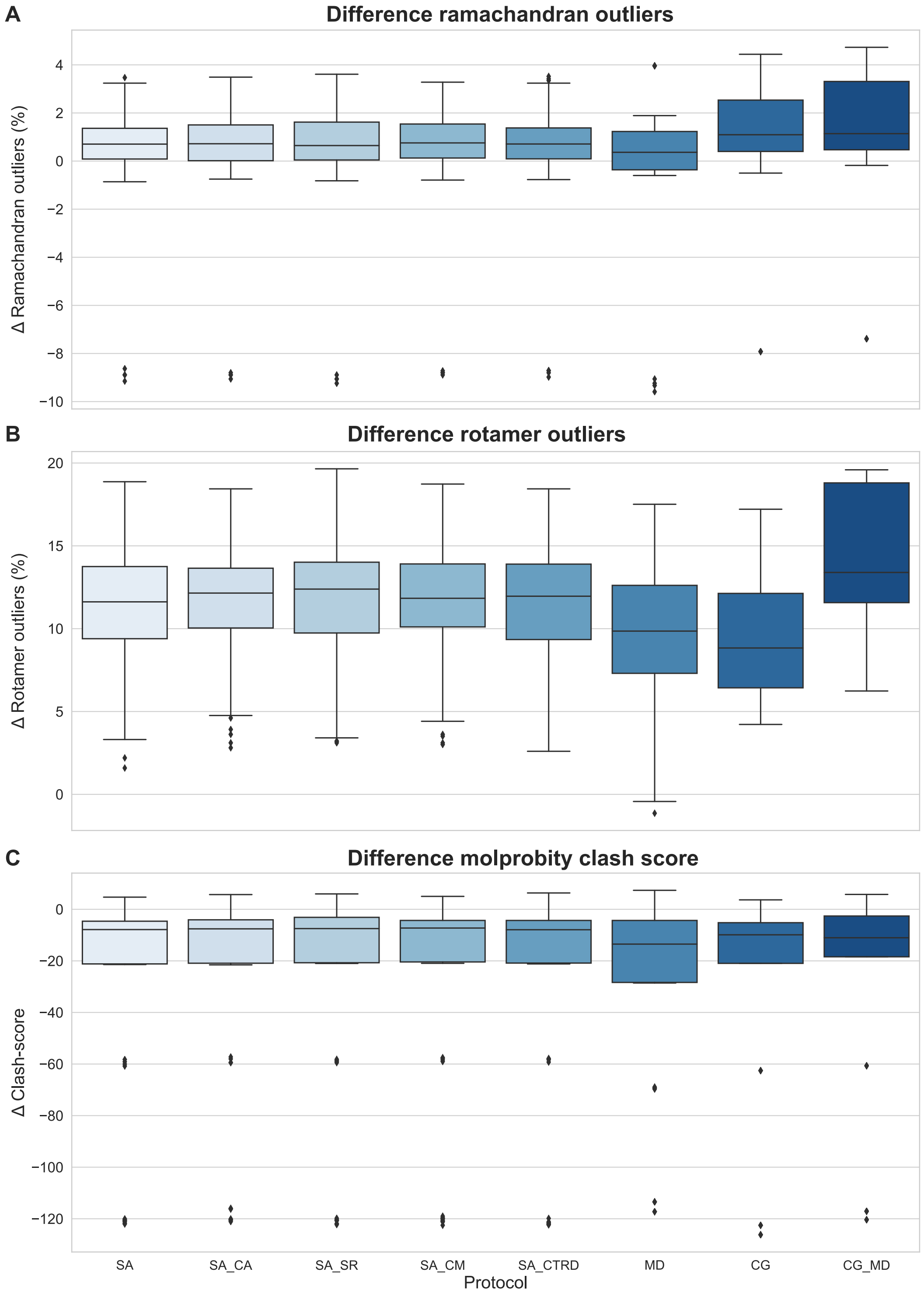

**Supplemental Figure 3.** Secondary structure analysis. The box plots represent delta values calculated by subtracting the reference from the model (model-reference) and are based on the top 4 refined models of all 14 complexes. Positive values indicate an increase with respect to the reference. **(A)** Delta helical residues are shown with respect to the reference structures. **(B)** Delta beta-sheet residues are shown with respect to the reference structures. **(C)** Delta turn residues are shown with respect to the reference structures. **(D)** Delta unstructured residues are shown with respect to the reference structures.

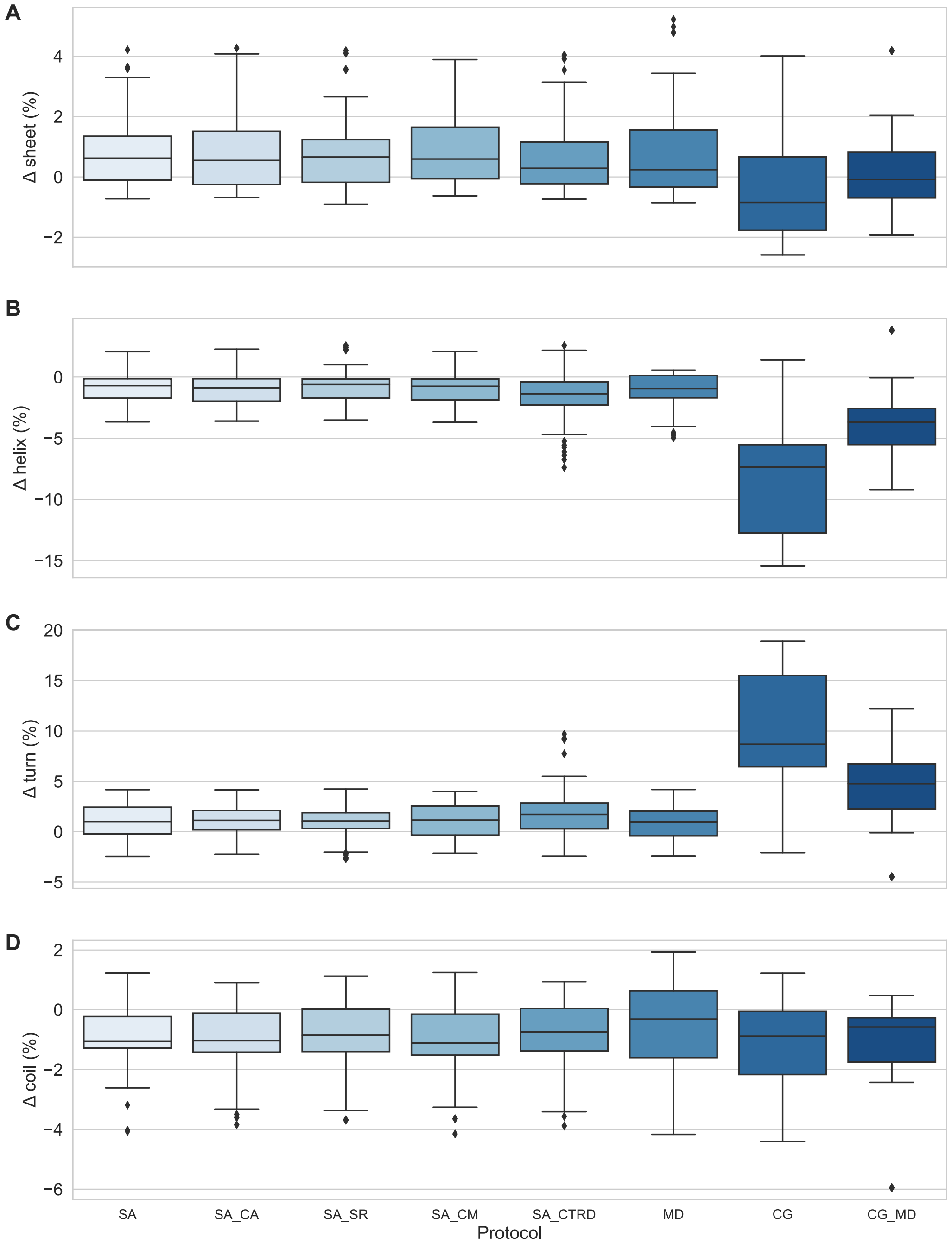

### SUPPLEMENTAL METHODS

#### Set up of an SA_CTRD Refinement Run

**Using the HADDOCK Webserver.** The interface refinement protocol using simulated annealing with centroid restraints can be accessed via https://haddock.science.uu.nl/haddock2.4/refinement which requires an account with an easy access level. Here, the full PDB file containing all chains is uploaded. In this file, modified amino acids, carbohydrates, ions and ligands should be formatted according to the HADDOCK2.4 library: https://wenmr.science.uu.nl/haddock2.4/library. When refining multiple starting conformations, an ensemble of structures should be provided. Next, *Simulated Annealing with Centroid Restraints* is selected from the dropdown menu. After pressing the next button, the numbers of uploaded models and each chain is presented. In the tab advanced sampling, multiple parameters can be tweaked. In order to run the basic protocol, no tweaking is necessary.

**Using a Local Computer.** To run the simulated annealing with centroid restraints refinement protocol using a local installation of HADDOCK2.4, each individual chain of a complex should be a separate PDB file. The various chains of a PDB file can easily be split using `pdb_splitchain.py` from pdb_tools. Here, modified amino acids, carbohydrates, ions and ligands should also be formatted according to the HADDOCK2.4 library: https://wenmr.science.uu.nl/haddock2.4/library. Next, unambiguous distance restraints should be generated if chain breaks are present, this can be done using the `restrain_bodies.py` script from haddock-tools (https://github.com/haddocking/haddock-tools). For ligands, unambiguous distance restraints must be generated using the `restrain_ligand.py` script from haddock-tools. After generation of the run directory, several parameters need to be altered in the `run.cns` file: `expand` (false => true), `randangle` (6 => 0) and `expansion` (0.4 => 0) (Table 3). This will result in an automatic generation of centroid restraints essential for the refinement. To ensure that the complexes are not docked during the rigid body phase, parameters `randorien` (true => false), `crossdock` (true => false), `ntrials` (5 = 1) and `rigidmini` (true => false) should be changed. These adjustments to the default parameters will result in protein complex refinement by simulated annealing with centroid restraints in HADDOCK 2.4. Other parameters that can be altered are `structures_0` (default = 1000), `structures_1` (default = 200), `anastruct_1` (default = 200) and `waterref` (default = 200), these are the number of generated models by HADDOCK. For refinement purposes we lowered all values to 50. If more input models were to be provided, the number of models to be generated can be increased to 100-200, for example, to limit the computational costs.

**Table 3:** Overview of the `run.cns` parameters to change in order to perform a HADDOCK interface refinement protocol using simulated annealing with centroid restraints.

| Parameter | Default | Interface refinement |
|---|---|---|
| **`Expand`** | false | true |
| **`randangle`** | 6 | 0 |
| **`expansion`** | 0.4 | 0 |
| **`randorien`** | true | false |
| **`crossdock`** | true | false |
| **`ntrails`** | 5 | 1 |
| **`rigidmini`** | true | false |
